# Interactions between *Brassica carinata* and its pollinators is shaped by managed beehives and neonicotinoid seed treatment

**DOI:** 10.1101/2024.11.30.626047

**Authors:** Henning Nottebrock, Shane Stiles, Jonathan G. Lundgren, Charles B. Fenster

## Abstract

*Brassica carinata* is a biofuel and animal feed crop with expanding global production. Although there is much research on common farming practices to improve yield, there is almost a complete absence of data on the dependency of yield through pollination services. Reciprocally, we lack information on whether *B. carinata* offers ecosystem services to pollinators. We observed almost 4000 pollinator visits, quantified different plant functional traits, including floral resources and examined the effect of supplementing fields with honey bee hives and the use of neonicotinoid seed treatment on seed yield and honey bee health. Data was collected from 35 0.404-ha sites with more than 800 focal *B. carinata* individuals across 2 years in the Prairie Coteau region of the Northern Great Plains. We found that pollinators (n = 28 species) are attracted to floral resources at different spatial scales. High visitation rates by pollinator species were associated with double the seed set in *B. carinata* relative to the lowest visitation rates. *Brassica carinata* adds floral resources to the agricultural ecosystem and therefore has the potential to increase pollinator health. However, species interactions are influenced by the use of insecticides and the presence of honey bees by managed beehives. In particular, insecticides alter the role of pollinators on crop pollination by reducing the positive impact of floral resources on pollinator-mediated yield and honey bee health.

## Introduction

Pollinators are a key component of global biodiversity and healthy ecosystems, providing vital ecosystem services including increasing crop and wild plant seed production (Kremen et al. 2007, Klein et al. 2007). However, of great concern is the threat to wild and domesticated pollinator health due to many factors including reduced habitat, lack of nutritional resources, and exposure to insecticides (Pollinator Health Task Force 2015, Kleijn et al. 2015, Giribaldi et al. 2021, Hanberry et al. 2021). Thus, loss of pollination services has important negative ecological and economic impacts that could significantly affect the maintenance of wild plant diversity, wider ecosystem stability, crop production, food security, and human welfare (Aizen et al. 2009, Potts et al. 2010, Millard et al. 2021, Murphy et al. 2022).

We have limited understanding of how the composition and the distribution of floral resources affects competitive and facilitative interactions mediated by pollinators in agricultural landscapes (Bottero et al. 2023, Cole et al. 2022, Labonté et al. 2023, Senapathi et al. 2017). Pollinators may also compete with individuals from the same species or with others for floral resources, depending on the overlap of co-flowering plants at different spatiotemporal scales (Nottebrock et al. 2017). For example, mass flowering crops that provide abundant floral resources such as our focal species *Brassica carinata*, might lower densities of pollinators in neighboring crops or nearby semi natural habitat (Holzschuh et al 2016). A challenge is to understand how pollinators contribute to maximize yield in agricultural landscapes at multiple spatial and temporal scales given the variable nature of the landscape, including different farming practices (Liss et al. 2013, Grab et al. 2017, Nottebrock et al. 2022). While there is increased evidence that improving nesting habitats and floral resources can have a positive effect on pollinator health in agricultural landscapes, including oilseed crops (Sponsler et al. 2023, Bottero et al. 2023), it is still not clear how wild and domesticated pollinators compete for floral resources and how farming practices, an important landscape level determinant of ecosystem composition, modulate plant-pollinator interactions at different spatiotemporal scales.

The demand for biofuels has a strong impact on plant-pollinator interactions in agricultural landscapes (Tudge et al. 2021). Oilseed field crops provide nectar and pollen resources for pollinators and can contribute to the maintenance of pollinator health when pesticides are absent (Stiles et al. 2021). Rotation and the planting composition of crop fields, as well as the availability of floral resources in the surrounding areas, are important farming practices affecting both the health of pollinators (Osterman et al. 2024) and the effectiveness of conservation agriculture in reversing the decline in pollinator health. An increasing number of studies demonstrate negative effects of pest control on pollinator health (Lundin et al. 2013, Mogren and Lundgren 2016), including on honey bee health and abundance (Lundgren 2017, Obregon et al. 2022). Thus, the spatiotemporal availability of contaminated resources and composition is important to understanding pollinator-mediated interactions in agricultural landscapes. However, there are almost no studies that shed light on the complexity of the interaction between wild pollinators, managed honey bees and insecticides within an agricultural ecosystem context. Although pollinator-mediated interactions have been an ecological research focus for decades (e.g., Fenster et al. 2004), there is no integrative mechanistic understanding of the complex interactions impacting pollinator health in agricultural ecosystems (Duncan et al 2015, Nottebrock et al. 2022). Thus, quantifying functional characteristics of plant species (plant traits) reveals ecosystem functioning directly mediating pollination services (Wood et al. 2015, Nottebrock et al. 2017, Vanbergen et al. 2013).

*Brassica carinata* has been identified as a biojet fuel feedstock with significant potential to contribute to USA defense needs and the sustainable and environmentally friendly fuel supply of the aviation industry (Sheeja et al. 2021). It is well adapted to the North Central US Midwest and in temperate Europe and has the potential to profitably lengthen crop rotations for area producers (especially as a winter annual crop in climates similar to the US southeast), and be cost competitive with petroleum and other biofuels. However, there is a complete absence of data on how much *B. carinata* oil crop yield relies on pollination services, or on the potential of *B. carinata* to supply ecosystem services to pollinators in the form of floral resources. Consequently, we use a trait-based approach (McGill et al 2006, Nottebrock et al 2017) to characterize plant traits that are correlated to the consumable floral resource (Kissling et al. 2012). We use a trait-based approach to quantify the effect of spatially and temporally distributed floral resources on pollinator health and crop yield to address four questions:

1. Does *B. carinata* rely on pollination services, and what are the pollinators?
2. How are carinata-pollinator interactions modulated by the spatiotemporal floral resource availability of *B. carinata* of treated field crop sites with or without insecticides and with or without the presence of honey beehives?
3. Do wild and honey bees compete for floral resources and have an additive effect on yield of *B. carinata*?
4. What are the consequences of *B. carinata*-pollinator interactions for ecosystem services using honey bee health as a metric?

## Materials and methods

### Descriptive biology of Brassica carinata

*Brassica carinata* (Brassicaceae), because of its high euric acid content (Falk 1999), has been identified as a new oil crop for jet fuel (Bouaid et al. 2005), with significant potential to contribute to the defense needs of the USA. Also known as Ethiopian mustard or carinata, it originated from a natural cross between *Brassica nigra* and *B. oleracea* followed by chromosome doubling (Prakash and Hinata 1980). Carinata is a self-compatible mass flowering, herbaceous, erect, highly branched annual with an extensively developed root system, closely resembling very robust rape seed with a height reaching over 2 m in some of our study sites. Carinata is increasingly being used as a winter annual cover crop, especially in the southeast United States (Kumar et al. 2020) and in the future may be an important component of the wheat/fallow rotation in the Upper Midwest. Carinata is currently produced commercially in South America and Australia. Carinata has a flower that is visited by many pollinating species (open, roughly radially symmetric, upright in orientation; Fenster et al. 2004, 2009) and shares a similar pollination system with canola. *B. carinata* has been reported to cross-pollinate 30% of the time (Velasco and Fernandez-Martinez 2009; Cheung et al. 2015), due to its flower structure and delayed anthesis (Cheung et al. 2015). However, few studies examine the effect of pollination services by insects to *B. carinata* (Stiles et al. 2021).

### Study design

We planted 36 0.404-ha (one acre) field sites of *B. carinata* in the Prairie Coteau region of eastern South Dakota representing various land-use intensities of agricultural landscapes of the Northern Great Plains. The Prairie Coteau is an approximately 52,000 km^2^ unglaciated region, 20% of which is uncultivated, resulting in a diverse mosaic landscape of agriculture and semi-natural habitat. We implemented a 2 × 2 factorial experimental design including *B. carinata* sites with and without honey bee hives and sites treated and not treated with neonicotinoid insecticide to better understand environmental effects on carinata-pollinator interactions. The 2 × 2 factorial design was implemented at each of 9 farms resulting in 36 field sites (Fig. 1 study design), but one site went missing due to conflicting agricultural usage, resulting into 35 field sites. We planted *B. carinata* on farming systems that had non-tilling and tilling systems The separation between tilling and non-tilling systems had no significance effect in preliminary models. Thus, to reduce the number of parameters in our models we dropped the till versus no till factor from our analyses. Instead, these effects of tilling practice were subsumed as a component of the random effect to our analyses (see section statistical analyses). Seed were purchased from an organic seed producer (GreenSeeds AG) and planted ∼19.8 kg/ ha. Half of the seed was treated with Poncho 6.68 mL/ kg, Red Colorant 0.21 mL/ kg, and water up to 0.18 mL/ kg. Poncho contains 2.26 kg clothianidin (a neonicotinoid compound) per 3.79 L. Following a hierarchical study design, we randomly selected within each field site five 1-m^2^ squares and five randomly marked focal plants within each square. We measured plant height, plant density, number of flowers, nectar volume and concentration at each time visited (details below). At the beginning of the growing season after plant establishment, every plot was evaluated to determine the overall density of the *B. carinata* individuals to calculate individual-based maps of each site.

**Figure 1.**
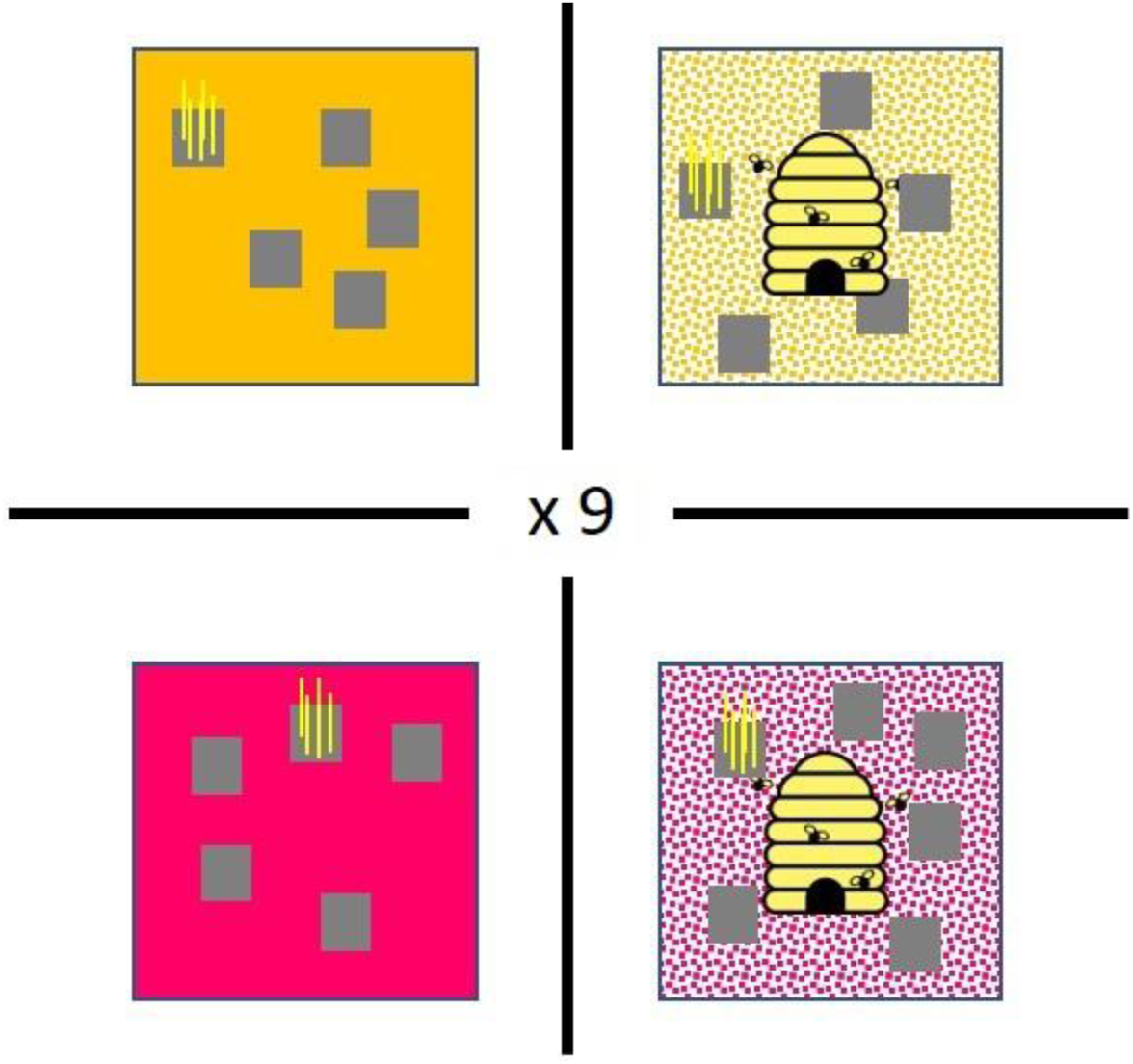
Factorial study sampling design to observe pollinators and to sample plant traits for the quantification of floral resources at plant, square and site scale. Colors of rose and yellow indicate sites treated with and without pesticides, respectively. The honey bee symbol shows the nested design of honey bee hive presents. Finally, the grey rectangles within the larger square and the yellow stripes show the nested block sampling design of 36 sites total (crossed sites of 18 seed treated sites and 18 presents of honey bee hives), 5 randomly selected 1m² squares (grey rectangle) and 5 randomly selected focal *B. carinata* individuals (yellow stripes) within each block. Finally, 35 (19 + 16) instead of 36 carinata sites were suitable for field work as 1 sites were destroyed through agricultural purposes.

### Pollinator visitation to Brassica carinata

To assess the role of pollinator dependent yield we excluded pollinators with pollinator exclusion bags and compared them to nearby, within 0.5 m, control plants of similar height and flower number. At each site we randomly located the pollinator exclusion treatment and controls throughout the 0.404-ha sites, with an average of two individual carinata plants bagged for pollinator exclusion and 10 other carinata plants to act as controls per site. At the end of the flowering season we located the plants and compared seed number per fruit for each treatment.

To assess the frequency and consequences of different pollinators on *B. carinata* yield, at each square, pollinator visitation was observed during peak pollinator activities (8am - 3pm). Pollinator observations at each square were conducted for 20 min. We noted the frequency of pollinator visits to each focal plant, identified to different groups of pollinator species based on average sizes (see Table 1 below), and made the distinction between visitations from a different plant and visitations within the same plant ranging from 1 up to 19 visitations per 20 min session and insect individual. Finally, at observational sessions heights of each focal plant, number of flowers on each focal plant, and number of flowers within the square were recorded. Nectar concentration was performed on focal plants with a refractometer (Bellingham and Stanley) and in addition, we noted the nectar volume and time of nectar sampling by squeezing the flower and creating categories of volumes (0.05, 0.01 and 0.1 mL) to estimate the sugar amount per flower and individual in gram by the formal brix% × volume nectar (see Stiles et al., 2021 for details). We then used the counted number of flowers per individual to calculate the respective sugar amount in gram per plant, square and site scale based on observed plant and flower densities in squares to create floral resource landscapes on small and larger scale that vary in available nectar sugar amounts for pollinators.

**Table 1.** Shows all observed pollinator visits to *B. carinata* during flowering. Pollinator names are at least identified to genus level. Pollinator frequency is the number of visits per pollinator species. Pollinator groups are common functional types of pollinators.

| Family | Species name | Pollinator frequency | Pollinator group (size) |
| --- | --- | --- | --- |
| Apidae | *Apis mellifera* | 2032 | Honey bee |
| Syrphidae | *Toxomerus marginatus* | 684 | Small hoverfly |
| Syrphidae | *Metasyrphus americanus* | 255 | Large hoverfly |
| Apidae | *Melissodes agilis* | 193 | Wild bee |
| Syrphidae | *Toxomerus geminatus* | 177 | Small hoverfly |
| Bombus | *Bombus spp.* | 91 | Bumble bee |
| Syrphidae | *Helophilus fasciatus* | 72 | Large hoverfly |
| Halictidae | *Lasioglossum anomalum* | 61 | Sweat bee |
| Cantheridae | *Chauliognathus pensylvanicus* | 55 | Soldier Beetle |
| Syrphidae | *Sphaeriohoria contigua* | 49 | Medium hoverfly |
| Syrphidae | *Syritta flaviventris* | 36 | Medium hoverfly |
| Pieridae | *Pieris rapae* | 33 | Cabbage white |
| Syrphida | *Meliscaeva cinctella* | 32 | Medium hoverfly |
| Halictidae | *Lasioglossum fuscipenne* | 24 | Sweat bee |
| Anthomyiidae | *Hydrophoria lancifer* | 19 | Calypttratae fly |
| Fanniidae | *Fannia canicularis* | 17 | Calypttratae fly |
| Sarcophagidae | *Sarcophaga johnsoni* | 10 | Calypttratae fly |
| Chloropidae | *Apallates spp.* | 10 | Grass fly |
| Apidae | *Anthophora spp.* | 9 | Wild bee |
| Nymphalidae | *Chlosyne gorgone* | 8 | Checkerspot |
| Muscidae | *Musca domestica* | 4 | Calypttratae fly |
| Tabanidae | *Tabanus spp.* | 3 | Horse fly |
| Calliphoridae | *Cynomya cadaverine* | 2 | Calypttratae fly |
| Tachinidae | *Ptilodexia rufipennis* | 2 | Calypttratae fly |
| Chrysomelidae | *Diabrotica barberi* | 2 | Corn root worm |
| Sarcophagidae | *Helicobia rapax* | 1 | Calypttratae fly |
| Sarcophagidae | *Ravina stimulans* | 1 | Calypttratae fly |
| Chrysomelidae | *Dibolia borealis* | 1 | Flea Beetle |

### Brassica carinata’s *seed set*

Once the fruits were mature at the end of the field season, we visited each site to harvest the squares and focal plants. The focal plants were harvested at ground level and placed into individual paper bags, while the remaining plants in the square were harvested and placed into a larger bag. All plants were dried for a minimum of ∼336 h at ∼60° C. Yield estimation was performed by harvesting the five focal plants contained within the 1 m^2^ squares. Each focal plant was weighed, the total fruits per focal plant were counted, and five fruits from each focal plant were selected to predict the total number of seeds per plant. Each fruit was weighed, and the number of viable and aborted seeds counted by verifying the presence of an endosperm in each seed. The prediction of yield on larger scales per field site was estimated through the extrapolation of biomass with seed number by measuring the biomass of the remaining plants in each square. Then we applied a regression of biomass and seed number and predicted yield at the site scale.

### Honey bee health

We placed four honey bee colonies with the common wooden hive system (supplied by Blue Dasher Farm, Estelline, South Dakota, USA, 57234) at half of the study sites with half of the sites planted with either treated with the neonicotinoid pesticide or with untreated seeds. Langstroth hives consisted of two deep boxes (24.45 cm deep) and shallow honey supers (14.61 cm deep) were added every 2 wk as needed. Hives were not provided supplemental feed nor treated for *Varroa* within the past 12 mo; colony weights per crop-management treatment at each farm were equivalent. The hives were weighed four times (approximately monthly, beginning in June) throughout the 2-yr growing season. We used the proportion of the colony weight by the number of hive boxes as the response variable to quantify the change of colony weight over the season.

### Statistical analyses

We used a trait-based approach to analyze carinata-pollinator interactions in agricultural ecosystems. We created predictor variables of nectar sugar amount at the plant, site and square scale to analyses the response variables of pollinator visits, seed set and honey bee hive health. Those floral rewards are computed instead of plant biomass and plant density because floral rewards are better predictors of plant-pollinator interactions (Nottebrock et al. 2017). Moreover, using a trait-based approach enables us to calculate predictor variables with sugar amounts contaminated with and without neonicotinoids directly affecting pollinators (Zioga et al. 2020). All three scales are representing important behavioral activities of pollinators reflecting mechanisms of pollination services (Burkle and Alarcon 2010). To relate sugar amounts to the response variables pollinator visitation and bee hive weight at different spatial scales, we calculated the available sugar amount at the day of floral reward observations. For the seed set model we calculated the average sugar availability at a day of floral reward observation weighted over the season. All statistical analyses were done with the software R (version 3.5.6) (R-project). We used the linear mixed effect models (GLMM) from the R package (lme4) (Bates et al. 2015). Response variables in two main models were a) pollinator visits (we assume Poisson errors and normal errors) and b) seed set. To ensure that pseudoreplication was not an issue we included categorical variables of site, square, date, time of day and species as random effects. In addition, we used linear mixed effect models to analyze the contribution of pollinators to carinata seed set and included a random slope of mean seed mass per fruit and as random intercept site, date and insect species. We used mixed models from the lme4 package to show differences of treatment effect on honey bee health by comparing the effect of treated sites and untreated sites to bee hive weight. We included site as random intercept and date as random slope to model a) and b). The variables of the resource availability were checked for correlation and we found only high correlation between two scales of our predictor variables that is site sugar and square sugar (r^2^ > 0.8). All other variables had low correlations of r^2^ < 0.6; thus, the variables are independent predictors for response variables (Dormann et al. 2009). However, to ensure comparability and avoid collinearity, variables of plant, square and site sugar were scaled and centered by the mean. Bee hive weight was analyzed with a linear regression model. To be able to include the hive weight change over time we included seasonal effects to the hive weight by using the proportion of weight change based on the hive weight before the season and each time when we visited the site and weight during the field season. We used a stepwise backward variable selection to compare best models with null models to identify the significance of each model by likelihood ratio tests (LRT) (Crawley 2012). Finally, the dependence of *B. carinata* to pollinators in our exclosure experiment was analyzed with an ANOVA allowing an unbalanced design. Finally, we calculated the relationship of *B. carinata’s* seed set per plant to pollinator visits of all our 20 min sessions per pollinator species and individual correcting for variation of different groups of pollinators, environmental variation of sites and variation of environmental conditions between sessions as random effects in an GLMM model.

## Results

### Brassica carinata *yield depends on pollinators*

*B. carinata* seed set depends on pollinators, with a significant difference between flowers that were closed to pollinators versus those that were open to pollinators based on ANOVA (F-test: 8.59 P < 0.005, Fig. 2, unbalanced design). Specifically, *B. carinata* seed set was more than doubled by high pollinator visitation rate relative to low visitation rate: there were min 1787 fertile seeds per plant (SE= 267.63) there was no pollinator visitation during our 20 min observation sessions, compared to max 4118 fertile (SE= 267.63) seeds per plant with high (18 visits, SE= 10.84) visitation rates (Likelihood-ratio-test of GLMM: χ²_df,1_ 30.32, P < 0.001, Fig. 3). Visitation rates represented the average number of visits to a plant over the 20-min surveys within the squares. Sites that included honey bee hives showed a significant higher number of honey bees visiting *B. carinata* plants (F-test= 4.00, P < 0.05, Fig. 4D, unbalanced design).

**Figure 2.**
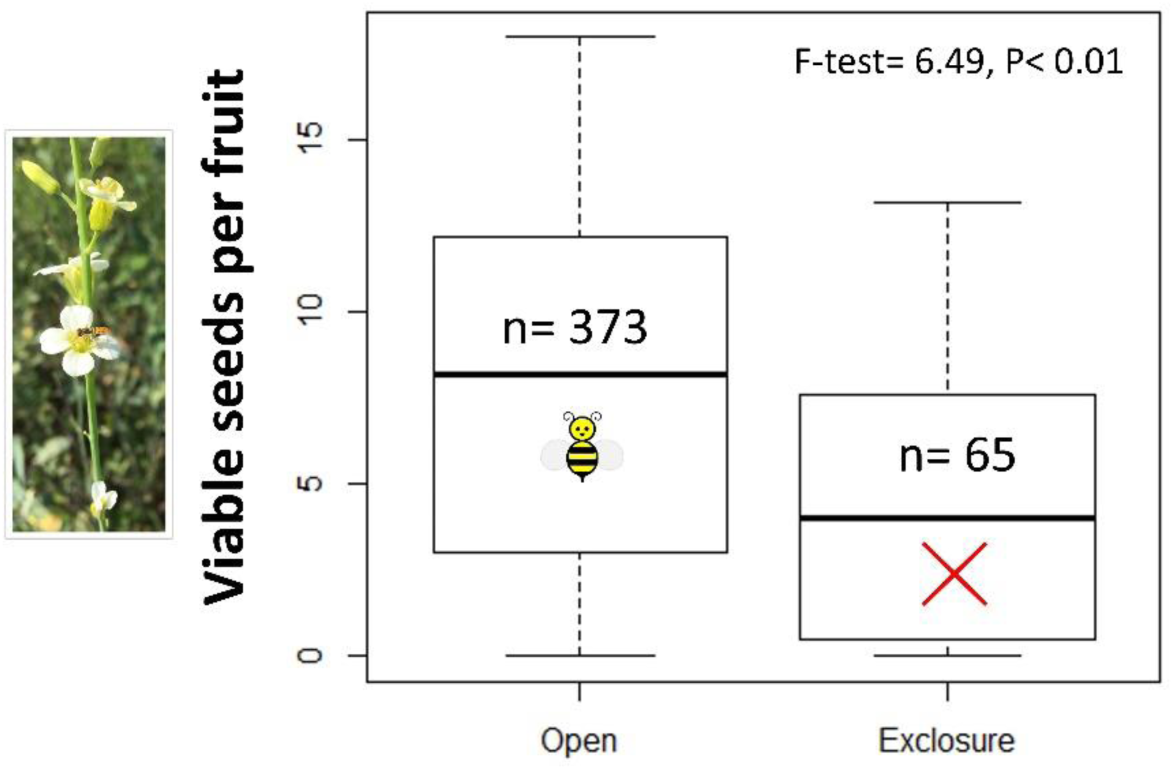
Experiment to quantify the importance of pollinators on reproductive success of *Brassica carinata*. We excluded pollinators (exclosure) from 65 individuals and used 373 individuals as reference with the possibility of being pollinated (open). Results show a significant difference between open and exclosure plants with fruits (p<0.01, F=6.49)

**Figure 3.**
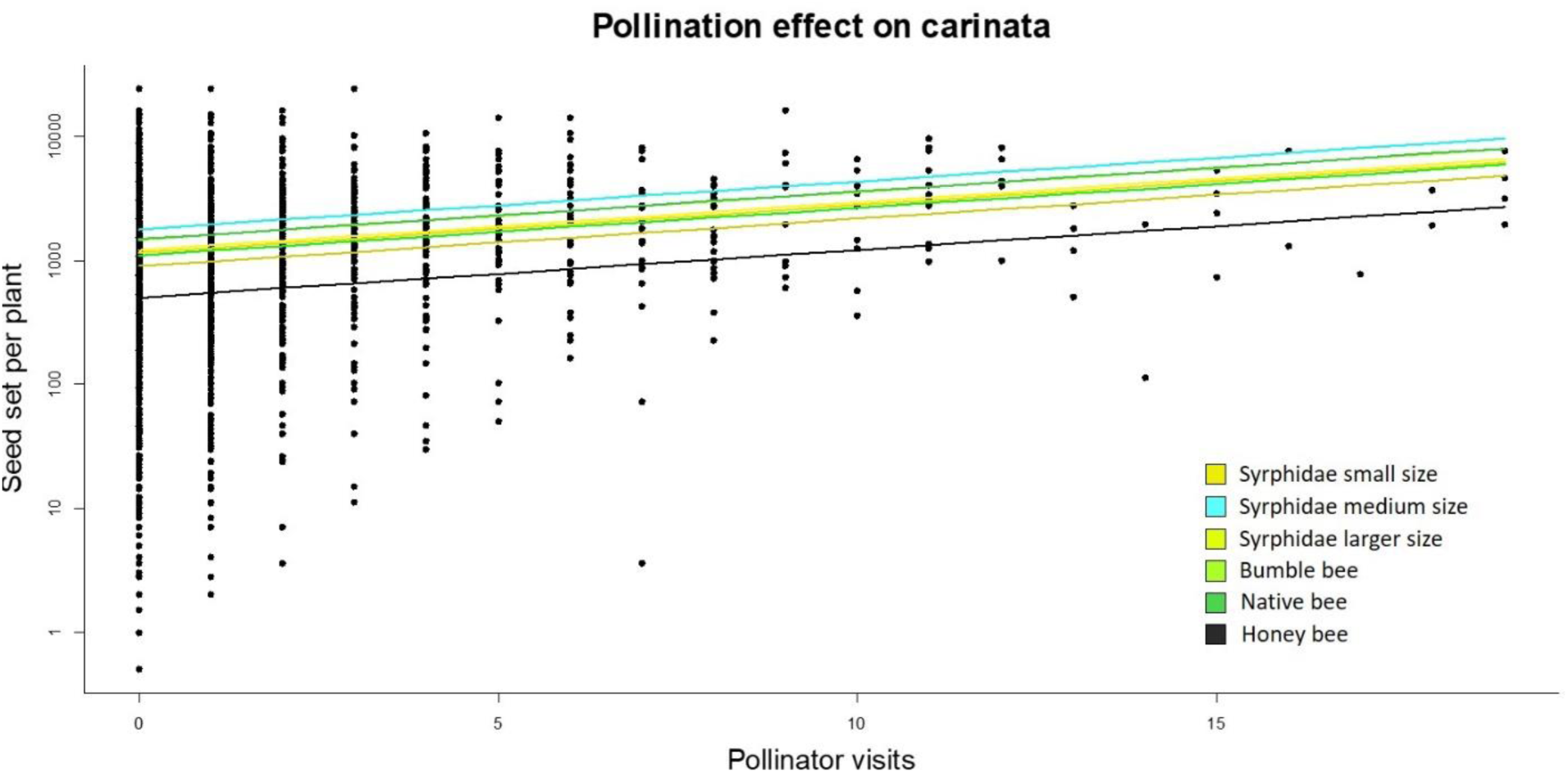
The effect of pollinators on *Brassica carinata’s* seed set with almost 4000 observations of pollinator visits and 800 focal plants. We quantified the effect of number of pollinator visits on focal seed set (to results: R²= 0.68, P< 0.0001). Focal plant individuals have been visited by 28 pollinator species (Table 1). Different colored lines indicate the most frequent groups of pollinators (see color legend for pollinator groups). The six groups account for 95 percent of all pollinator visits. 2209 pollinator visits when bee hives are present and 1750 without bee hives present. Please note that y-axis is log transformed. Pollinator names, frequency and groups are in Table. 1.

**Figure 4.**
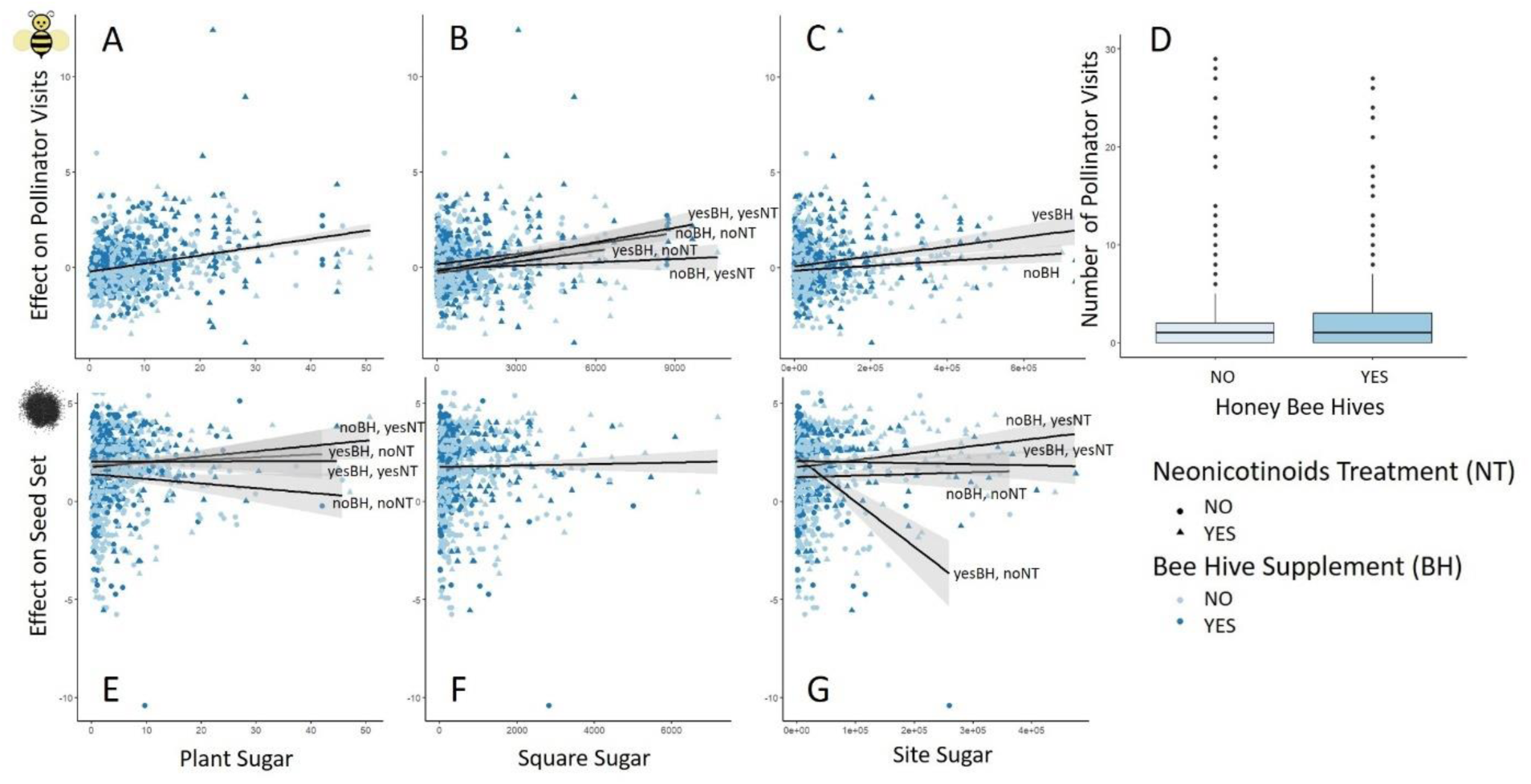
Positive or negative effects of floral resources at different spatial scales on pollinator visitations of B. carinata A to C, on B. carinata’s seed set E to F including the interactions with the supplement of honey bee hives and see treatment. Effect sizes are calculated as the estimated marginal means (emms) and the standard error (SE) of the corresponding model (see results) performed with the ‘emmeans’ package in R. Floral resource availability is represented at a plant scale, at a neighborhood scale (square) and at a site scale. Treatments are seed treatment with neonicotinoids and sites with honey beehives blue. Effects that are not represented in the minimal adequate model are not shown. The marginal and conditional R² for the pollinator visitation rate model is 0.10 and 0.82, respectively and for the seed set model 0.21 and 0.86. Units of nectar sugar are shown in [g].

### Nectar sugar resources modulate carinata-pollinator interactions depending on spatial scale

At the plant scale, sugar amount was positively associated with pollinator visitation (χ²_df,1_: 97.09, P < 0.001, Fig. 4A). Examining sugar amount at the square scale showed higher visitations of pollinators with more sugar amounts for all treatments with or without honey bee hive present, and with or without neonicotinoid seed treatment (χ²_df,1_: 21.42, P < 0.001, Fig. 4B). At the site scale, we found that pollinator visits increased with increasing nectar sugar availability, especially on sites where honey bee hives were present (χ²_df,1_: 14.20: < 0.0001, Fig. 4C). Pollinator visitation rates were lower on sites treated with neonicotinoids, and on sites with honey bee hives, there was significantly higher number of pollinator visits (χ²_df,1_: 5.71, P < 0.05, Fig. 4D).

Effects of nectar sugar resources on *B. carinata* seed set at the plant, square and site levels: At the plant scale, all three interactions of plant sugar, honey bee hive supplement and seed treatment are significant (χ²_df,1_: 96.90, P < 0.0001, Fig. 4E). We find that plant sugar availability attracts pollinator visits, which results in an increase *of B. carinata* seed set especially for treated sites without honey bee hive presence. However, this effect turns into a negative effect when *B. carinata* plants are non-treated with neonicotinoids and honey bee hives are not present. When honey bee hives are present with or without treated pesticide seeds we detect almost no effect of plant sugar on seed set. At the square scale, we find a slight positive effect of nectar sugar availability on plant seed set over all possible interactions (χ²_df,1_: 12.61, P < 0.001, Fig. 4F). At the site scale, we find different effects of all possible interactions being significant (χ²_df,1_: 176.38, P < 0.0001, Fig. 4G). We find a positive effect of site nectar sugar availability on plant seed set especially when sites are treated and no honey bee hives are present. This effect turns into a slight negative effect when honey bee hives are present and sites are treated with pesticides. Seed set is dramatically decreased at the site scale by the addition of honey bee hives without seeds being treated with neonicotinoids. Where honey bee hives are absent and seeds are not treated with neonicotinoids then the effect of site sugar availability at the site level turns into a slight positive effect on seed set consistent with aforementioned results indicating the positive role of pollinators in carinata yield.

### Honey bee health

We observe a positive effect of nectar sugar availability at the plant scale and a negative effect of site sugar availability on the weight of honey bee hives (χ²df,1: 5.22, P < 0.001, Fig. 5A and 5B). We found a clear signal of a negative effect on bee hive health, based on hive weight change over time on treated sites (F = 4.34, P = 0.04, Fig. 5C).

**Figure 5.**
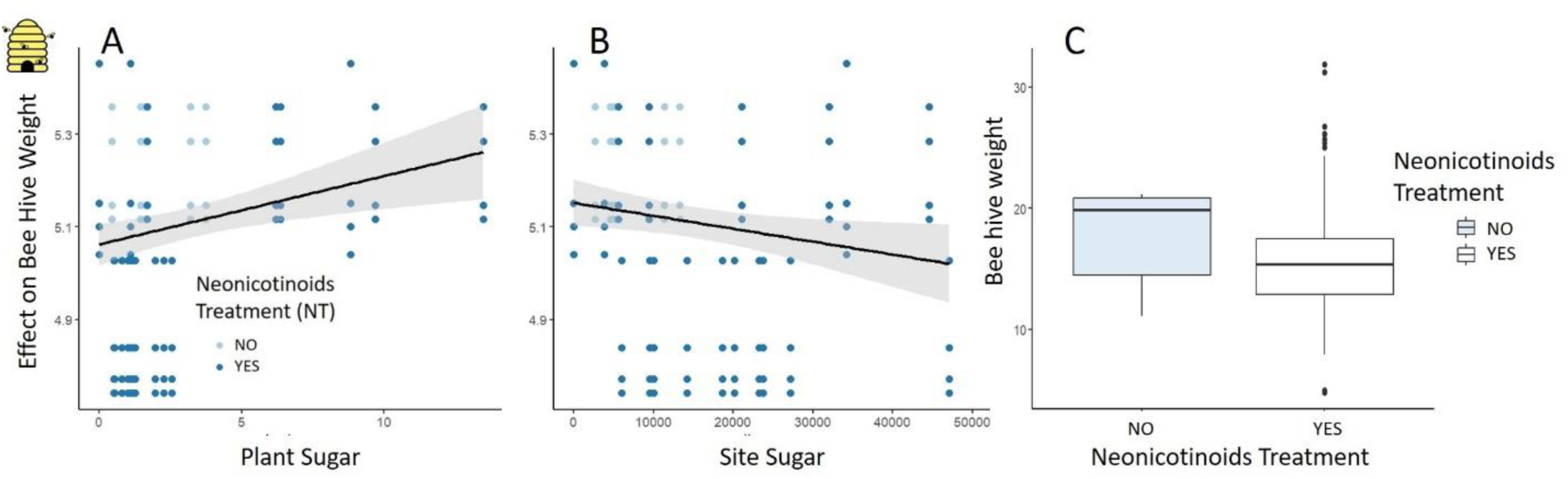
Positive or negative effects of floral resources at different spatial scales honey bee health **A and B.** Treatments are seed treatment with neonicotinoids shown in **C**. Effect sizes are calculated as the estimated marginal means (emms) and the standard deviation (sd) of the corresponding model (see results) performed with the ‘emmeans’ package in R. Effects that are not represented in the minimal adequate model are not shown. R² for the linear model is: 0.49. Units of nectar sugar are shown in [g].

## Discussion

We demonstrate that *B. carinata* yield is strongly dependent on pollinator services in the Prairie Coteau region of South Dakota. We also show that pollinator attraction to *B. carinata* is determined by the availability of floral resource at both small (plant, square) and large (site) scales. However, interactions between *B. carinata* and pollinators, especially honey bees interacting with wild bees, is strongly context dependent on the scale of the interaction and whether the field is treated with neonicotinoid pesticides and/or whether honey bee hives are present. Below we discuss these points in turn and conclude by noting important future research goals that have policy implications.

### Brassica carinata yield reliance on pollination services

When pollinators are abundant, their presence can double seed *set for B. carinata*. The positive effect of honey bee frequency on seed set might be biased by the presence of abundant honey bees both because of our hive supplementation treatment, also because there are so many summer hives in the region (Holzschuh et al. 2016, Giribaldi et al. 2013). Of the wild pollinators, medium sized syrphidiae have the largest effect on pollination success. Our research adds to the increasing number of studies that demonstrate the importance of syrphidae for pollination service (Dyle et al 2020, Földesi et al 2021).

Self and between plant pollination without pollinators will be dependent on wind and schedules of anther dehiscence and stigma receptivity. However, a diverse community of pollinators likely augments abiotic pollen transfer and do so throughout the day, thus increasing the likelihood of successful pollination and seed production. Our results are perhaps not surprising given the similarities between carinata and canola. Canola is typically pollen limited, i.e., yield is limited by inadequate transfer of pollen but yield increases with increasing density of bees (Morandin and Winston 2005). Parallel findings have been found with syrphid fly pollination of rapeseed (*B. napus*, Jauker and Wolters 2008; Haenke et al. 2014). In all studies on canola conducted throughout the world (e.g., Brazil, Europe, Canada, Australasia) wild and introduced pollinator service is associated with increased yield in terms of fruit-set, seed number, and even oil content (e.g., Sabbahi et al., 2005, Free and Nuttall, 1968, Klein et al. 2012) resulting in important economic benefits to the producer, e.g., increasing market value to the producer of up to 20% in Sweden (Bommerco et al. 2012). The contribution of pollinators to canola yield varies by variety (Hudewenz et al. 2014, Garratt et al 2018), thus it is imperative that future pollinator studies consider varietal differences in floral morphology, and floral traits for *B. carinata*.

### Effect of neonicotinoids on pollinator visits to carinata and on carinata yield

Neonictinoid insecticide is known to modulate pollinator behavior of wild and honey bees and is likely contributing to the pattern of pollinator visitations observed at our sites (Rundlöf et al. 2015). However, the abundance of honey and wild bees are not directly affected by coated seeds based on visitation rates quantified at the level of the squares, likely because ingestion of the pesticide reduces bee movement (Wintermantel et al. 2022). Low levels of neonicotinoids found in the nectar and pollen of plants is sufficient to deliver neuroactive levels to their site of action, the bee brain (Moffat et al. 2015), which we suggest leads to reduced movement (Moffat et al 2016). Thus, the contribution of honey bees to seed set at the site level is lower when the site has been treated with neonicotinoids. While the presence of honey bee hives increases the importance of honey bees for pollination visits on neonicotinoid treated sites, the effect of nectar sugar at different spatial scales remains negative on seed set in neonicotinoid treated sites, especially for honey bees at the site scale.

We primarily focused on the interplay between farming practices, floral resources of carinata, and pollinator visits, and found that wild pollinator visits generally increase with increasing floral resources at plant and site scales (Fig. 4A - C). When honey bee hives are absent and seeds are treated with neonicotinoids then the increase visitation of wild pollinators with larger carinata floral resources is similarly reflected in higher seed set at the same scales (Fig. 4 E - G) emphasizing the importance of wild pollinator services for *B. carinata* yield. Adding honey bee hives likely increases competition for floral resources between honey bees and wild pollinators which may result in lower pollination success since honey bees generally are less effective pollinators than wild pollinators (Osterman et al. 2023).

In a previous study (Stiles et al. 2021), where we included landscape scale attributes, we found that the heterogeneity of surroundings to the focal carinata sites was significantly positively associated with pollinator species diversity which were in turn significantly positively associated with carinata yield. Thus, we may conclude that the spatiotemporal availability of mass flowering crops such as in our study influences pollination services in competition between honey bees and wild pollinators. Landscape elements as flowering strips and semi natural habitats that support wild bees may also contribute to higher yields when farming practices do not disrupt the interplay of honey bees and wild pollinators.

### Scale effects: plant and pollinator interactions

Our study provides a comprehensive understanding of the factors influencing pollinator services and crop yield at multiple scales. The importance of honey bees to carinata yield is reduced as the amount of floral resources for wild pollinators increases. While small-scale plant-pollinator interactions between honey bees and wild bees increase yield of *B. carinata* observed at the square scale, this positive synergy is reduced when plants have been treated with neonicotinoids. Nevertheless, this effect where the higher frequency of honey bees may compensate the effect of neonicotinoids on pollinators (Lundin et al 2015), does not result in higher seed set with increased honey bee visits on sites where bee hives are present. Perhaps honey bee visitation rate is higher but even less effective for pollination services because the movement is restricted and thus the saturation effect is stronger due to lesser directed flights (Desneux et al., 2007; Hopwood et al., 2012; van der Sluijs et al., 2013; Feltham et al., 2014;). However, this needs to be confirmed on a landscape scale if the surroundings are intensely treated with pesticides and there is a comprehensive lack of nutritional resources.

In contrast at the smallest scale, local abundance of plant nectar sugar acts is a compelling magnet for pollinators, intensifying visitation rates and positively influencing seed set for both wild and honey bees (Moeller 2004, Albrecht et al 2012). However, the impact of neonicotinoid pesticides on these dynamics is complex. While plant sugar lures more pollinators, the presence of pesticides alters the behavioral patterns of both wild pollinators and honey bees, potentially restricting their movement despite sugar availability at the larger scale. This interplay underscores the balance between attracting pollinators and mitigating their potential exposure to harmful substances such as neonicotinoids or other pesticides. This is especially concerning for the health of honey bee populations, which exhibit a pronounced reliance on plant sugar. Moreover, pollination services at the larger site scale shows that higher frequencies of honey bees decrease pollinator visits possibly due to competition with other pollinators (Osterman et al 2021), reflected in the lowering effect of honey bees on *B. carinata* seed set in comparison with wild pollinator effects. This result highlights the importance of wild pollinators to assure *B. carinata* yields, even when the honey bees increase seed set, wild pollinators appear to play a major role on carinata seed set.

### Effect of neonicotinoids on honey bee hive health

Honey bee health increases with *B. carinata* floral resource availability. However, we found that honey bee health decreases when these floral resources are contaminated by the pesticide, especially at larger scales. Honey bee hive weight also decreases in oil seed rape fields treated with neonicotinoid pesticides (Wintermantel et al. 2020). Thus, treatment of carinata fields with neonicotinoid pesticides both reduces the positive effect of pollinators on yield and reduces hive health. Our findings highlight the critical need to consider the impact of pesticide-contaminated floral resources in understanding and optimizing pollination services in *B. carinata*. The interplay between contaminated resources and pollinator behavior emphasizes the importance of managing chemical exposure to maintain effective pollination and crop yield.

### Conclusion

While honey bees contribute to increased *B. carinata* yield, wild pollinators play a pivotal role at larger site scales, particularly in enhancing yield regardless of neonicotinoid use. However, the presence of neonicotinoids significantly reduces honey bee health, impacting their long-term pollination effectiveness. This underscores the critical importance of wild pollinators in ensuring optimal yield and highlights the complex interactions between chemical exposure and pollinator behavior. Therefore, understanding the multifaceted interplay between honey bees, wild pollinators, plant sugars, and pesticide exposure is paramount for conserving biodiversity and safeguarding essential pollination processes (Bottero et al. 2023). To optimize crop yields and ensure pollinator health, policies should encourage farming practices that protect wild pollinators, reduce neonicotinoid exposure, and support habitat conservation (Stiles et al. 2021). Farmers can benefit from maintaining wild pollinator populations, as they are crucial for sustained productivity at larger scales, even in the presence of honey bee hives.

## Acknowledgements

We thank N. Petersen, J. Gelderman, and J. Smithers for their assistance in the field and laboratory. We thank the farm producers who allowed us to work on their properties. Additionally, we thank Dimitry Wintermantel for helpful comments on a former manuscript version. This research was funded by the North Central Sun Grant Initiative (USDA/DOE) SA1500640 and the Oak Lake Field Station. Funding comes also from HORIZON EU RestPoll project ID 101082102.

